## Supplemental Figures for "Molecular development of muscle spindle and Golgi tendon organ sensory afferents revealed by single proprioceptor transcriptome analysis"

Supplemental Figure 1: Analysis of *Rx3:FlpO* mice

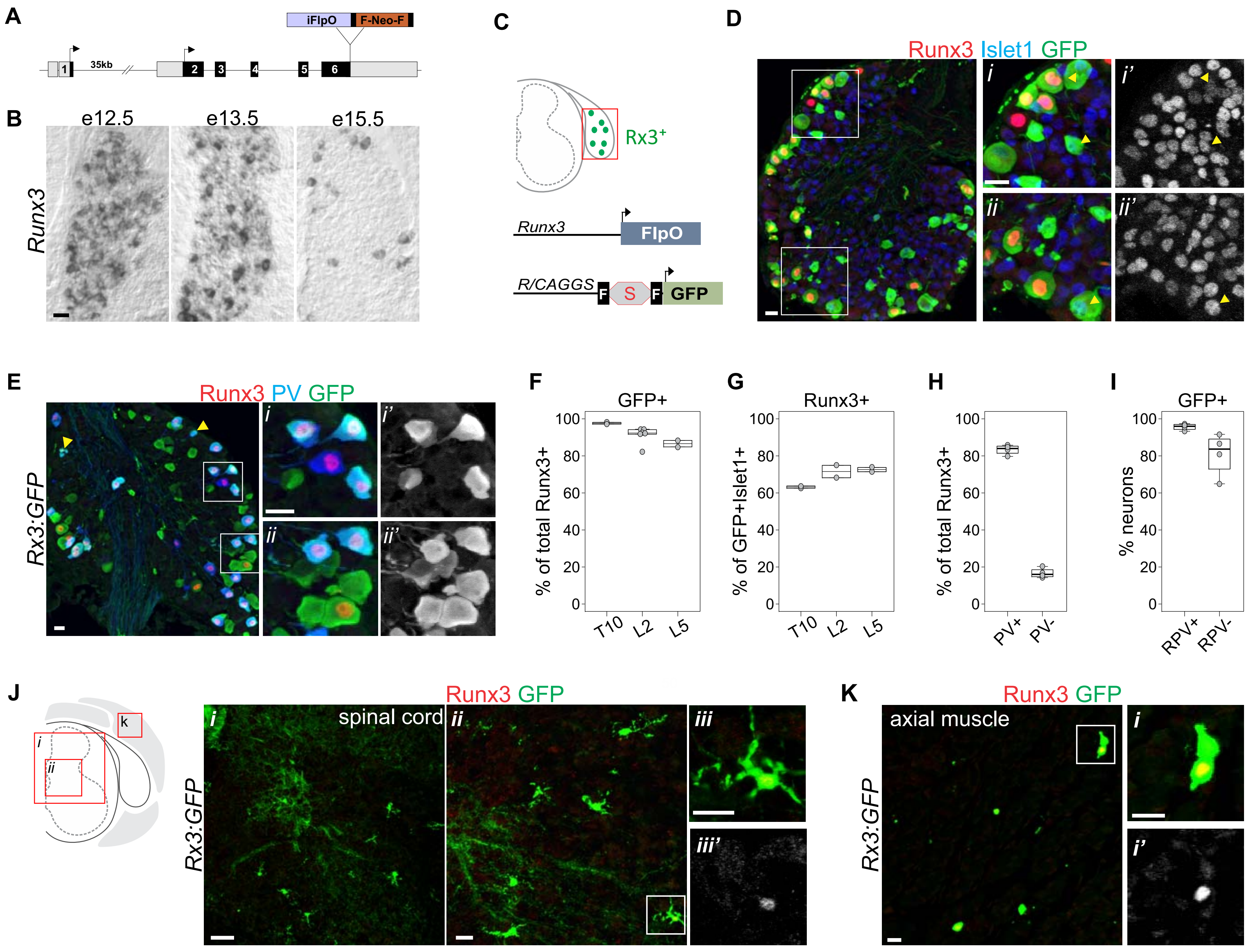

Supplemental Figure 2: Analysis of *PVRx3:tdTomato* mice

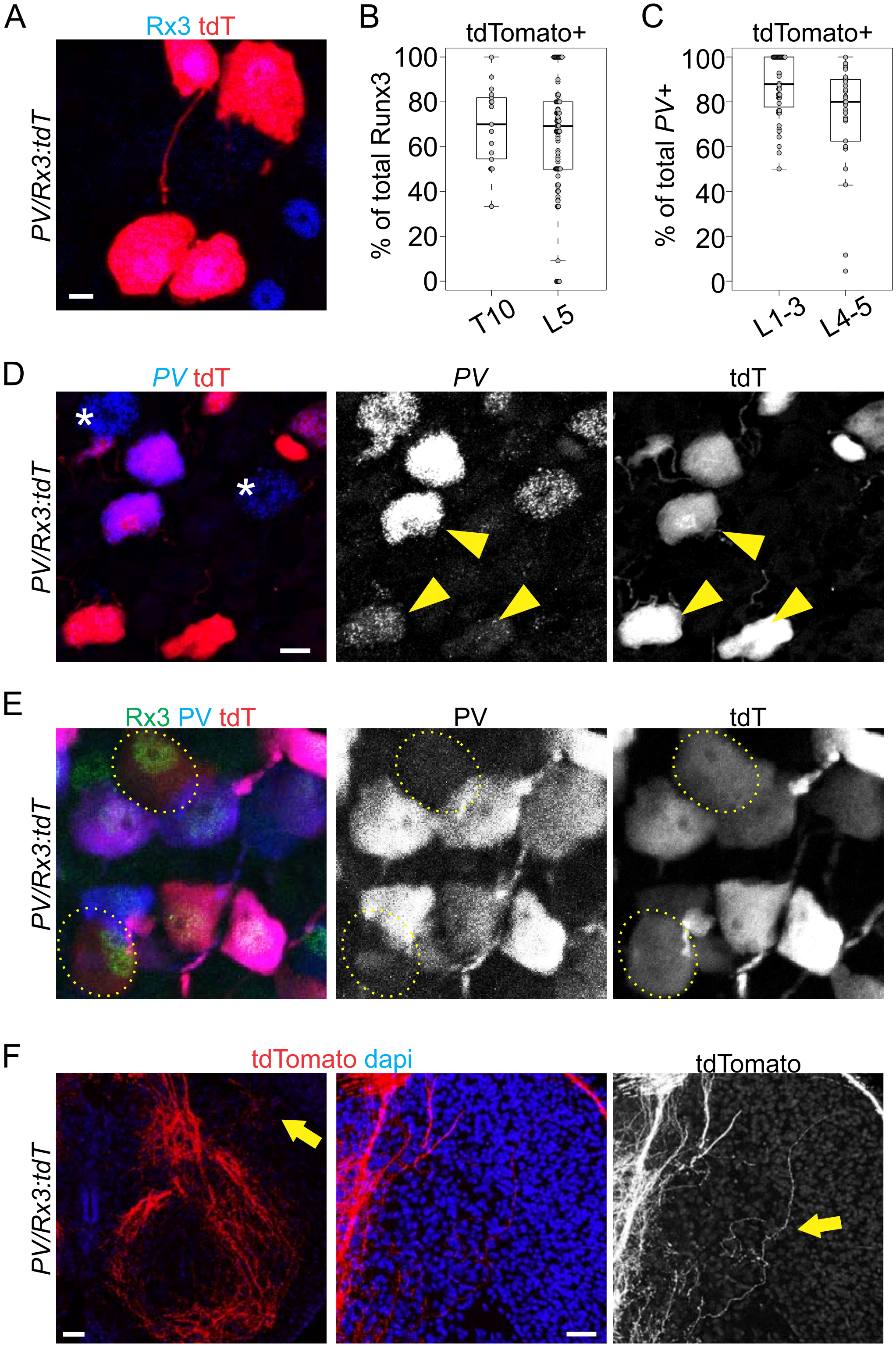

Supplemental Figure 3: Analysis of *PV/Rx3:tdTomato* cutaneous and interosseous membrane afferents

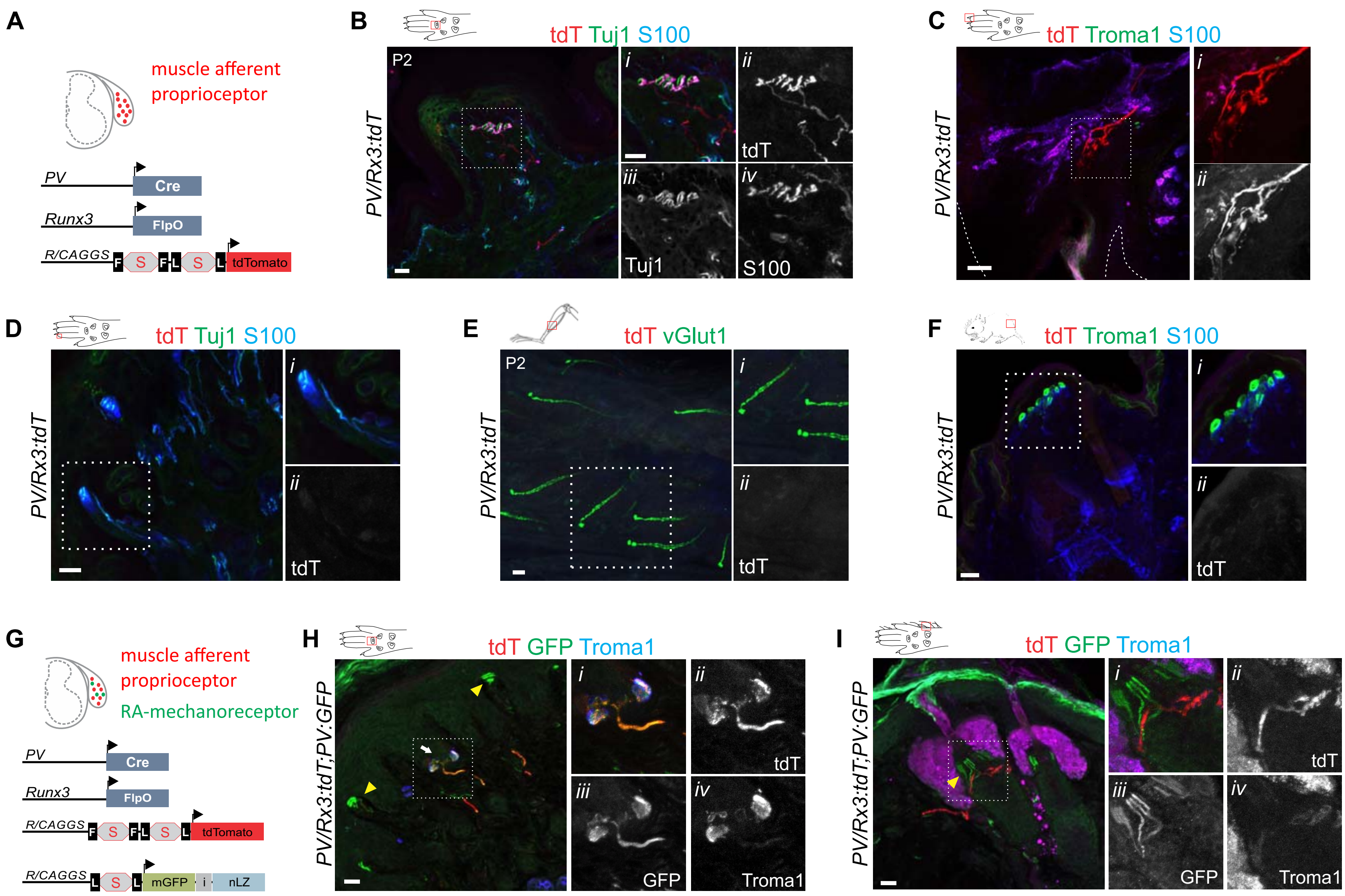

Supplemental Figure S4

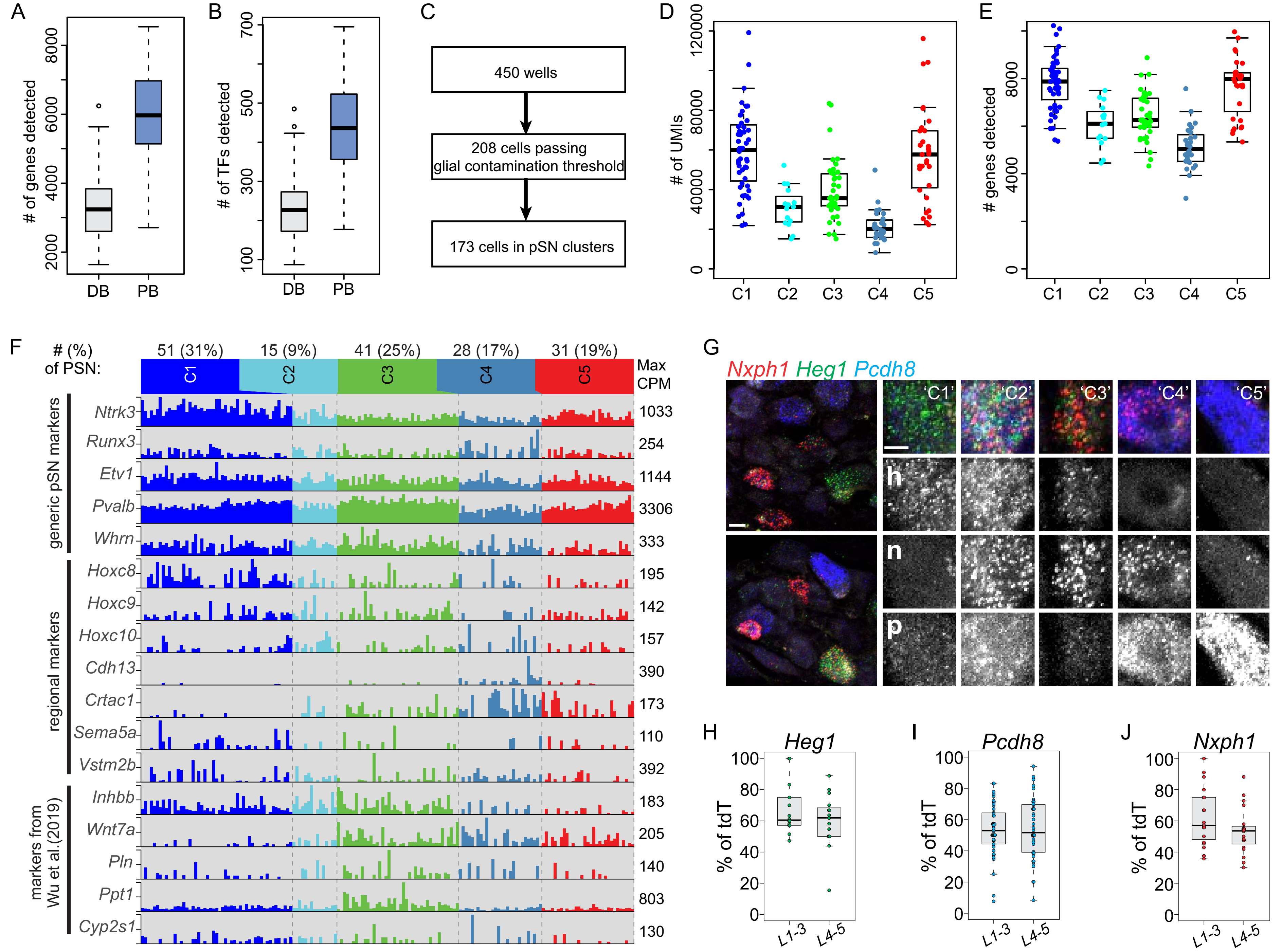

Supplemental Figure S5

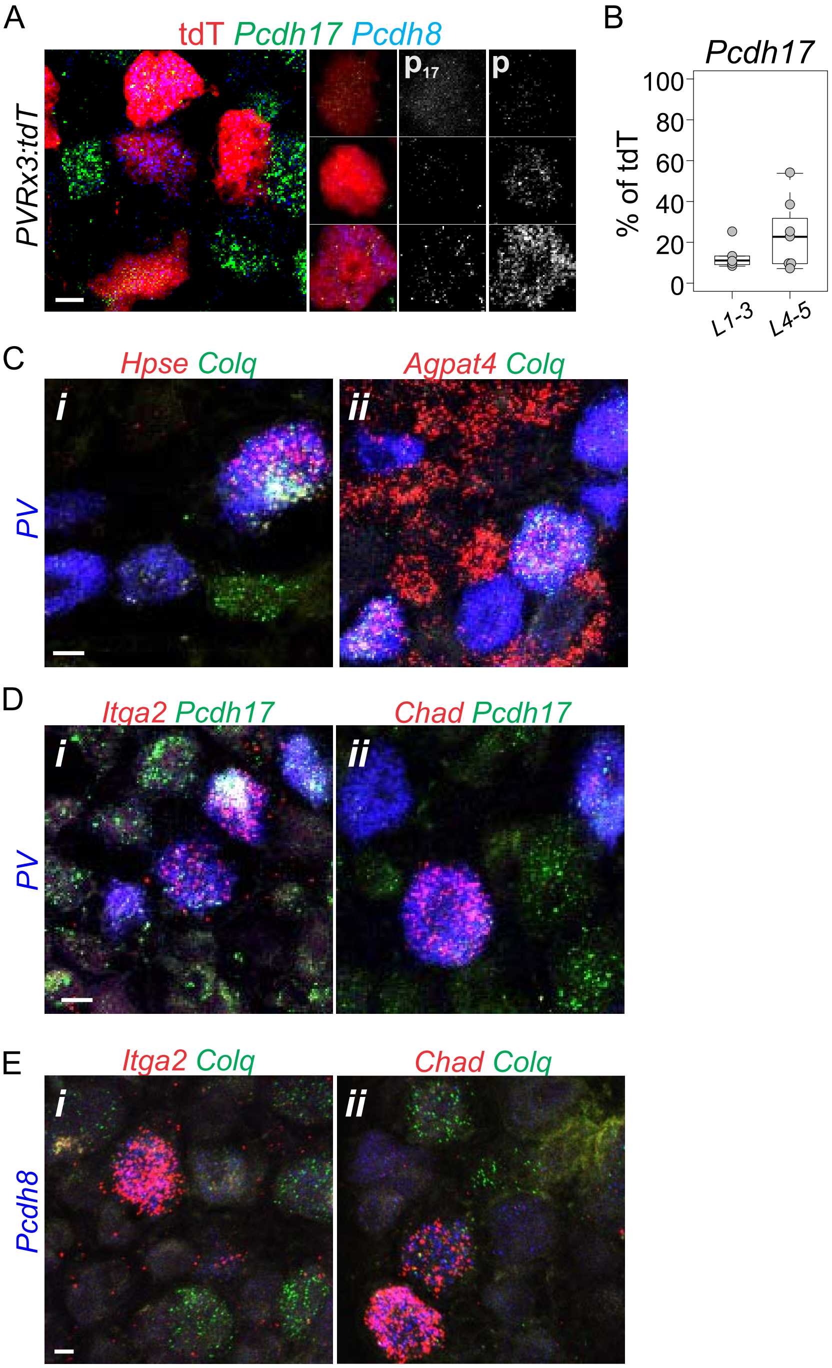

Supplemental Figure S6

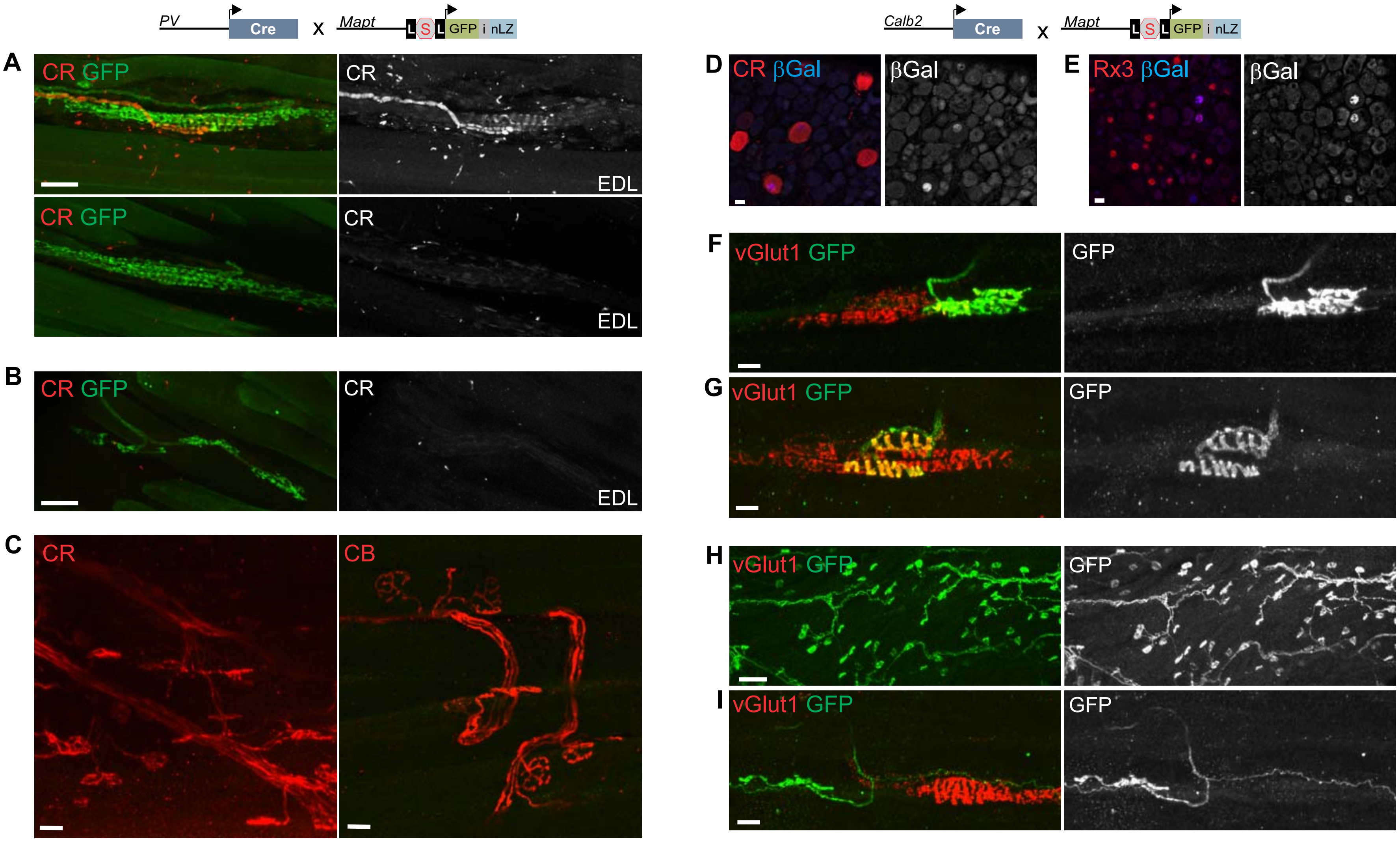

Supplemental Figure 7

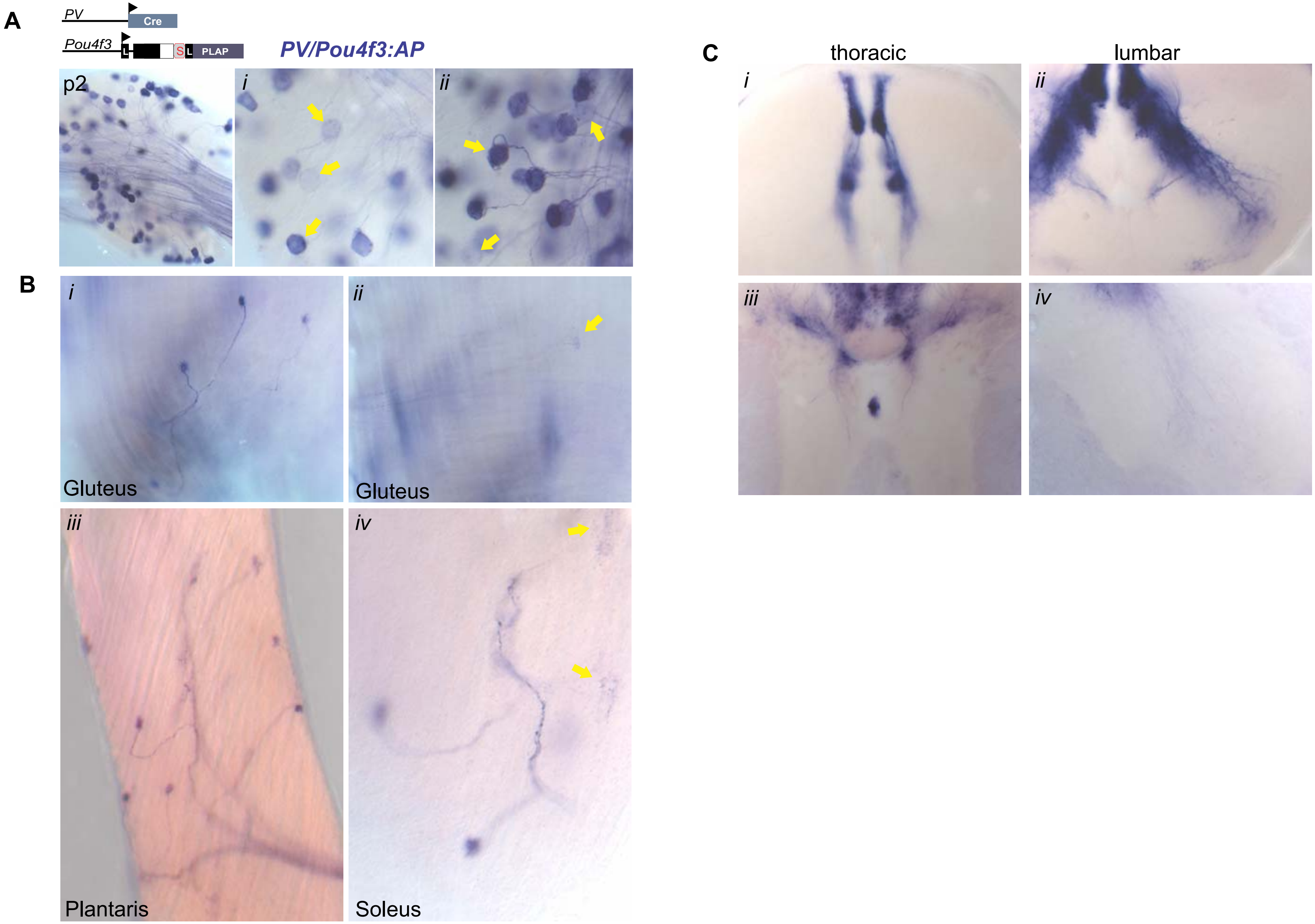

Supplemental Figure S8

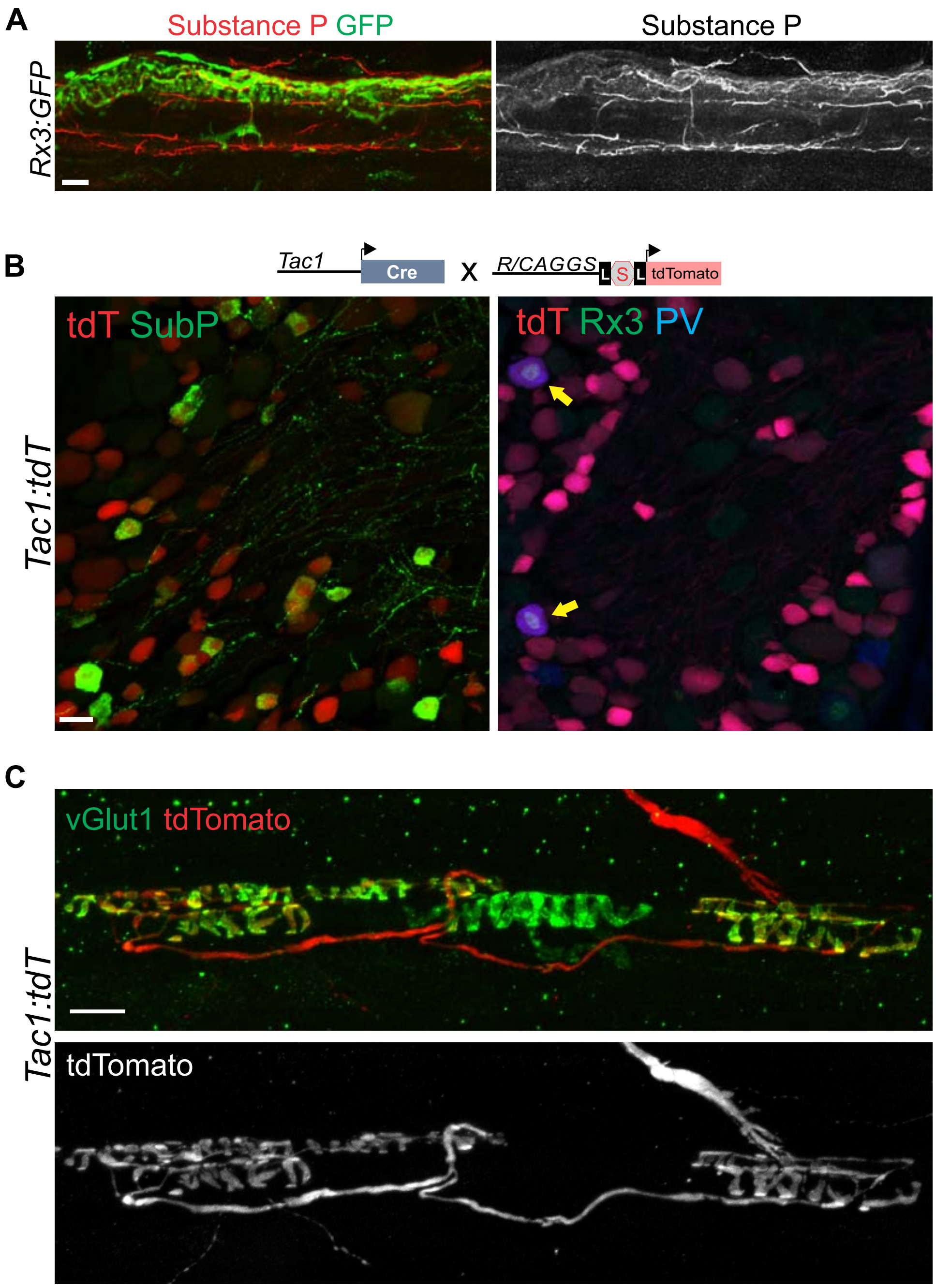

### Supplemental Figure S9

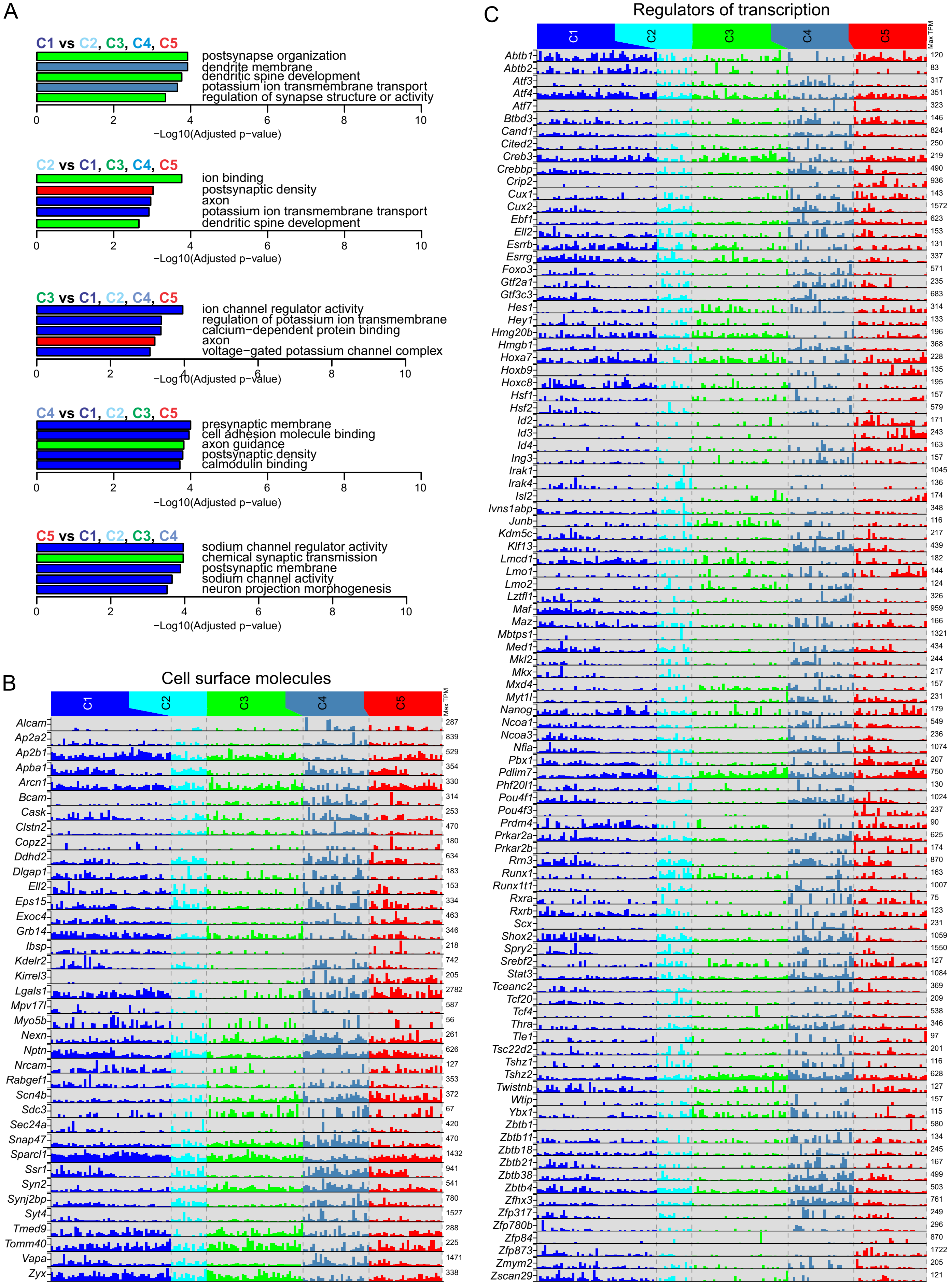

### Supplemental Figure S10

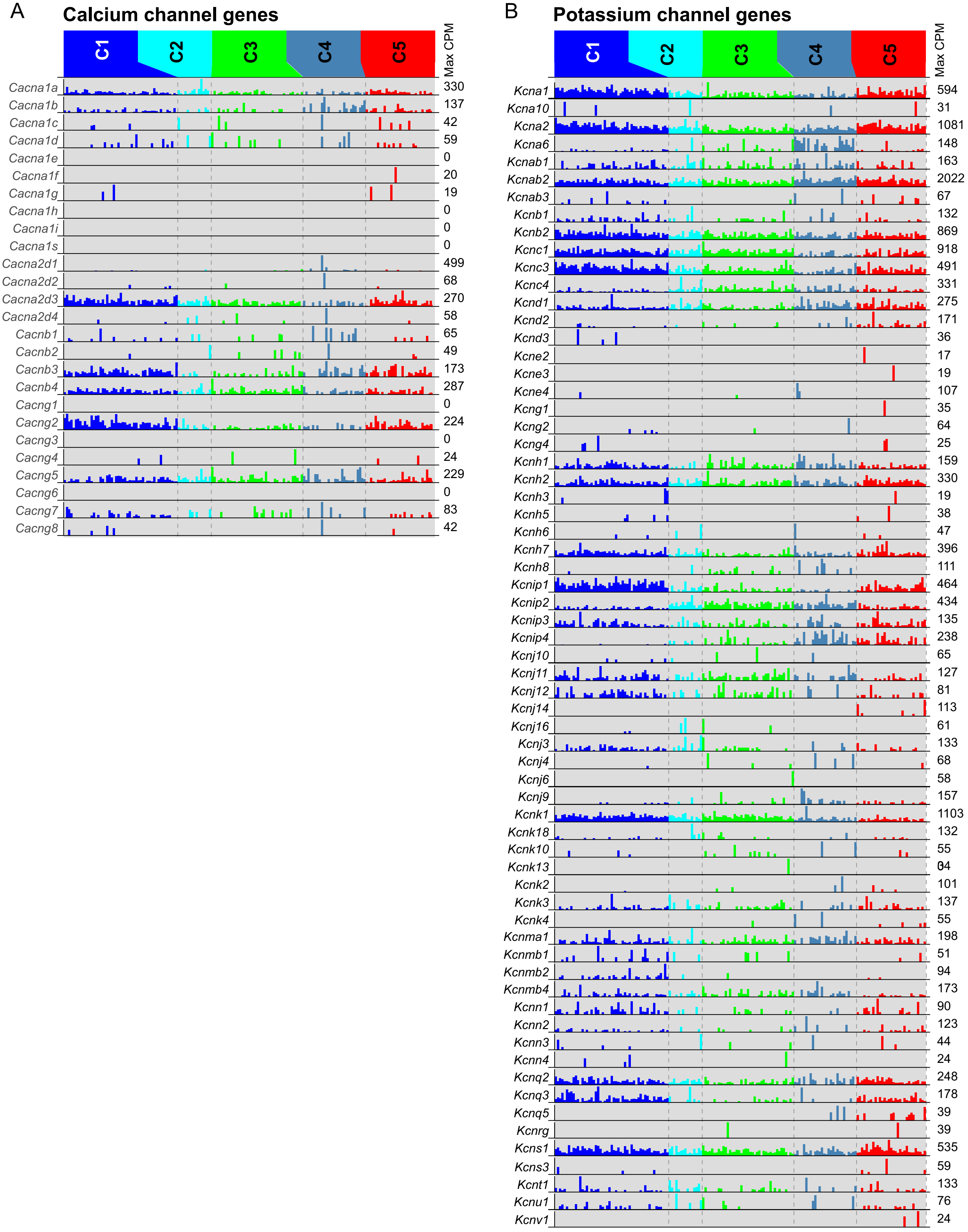

Supplemental Figure S11.

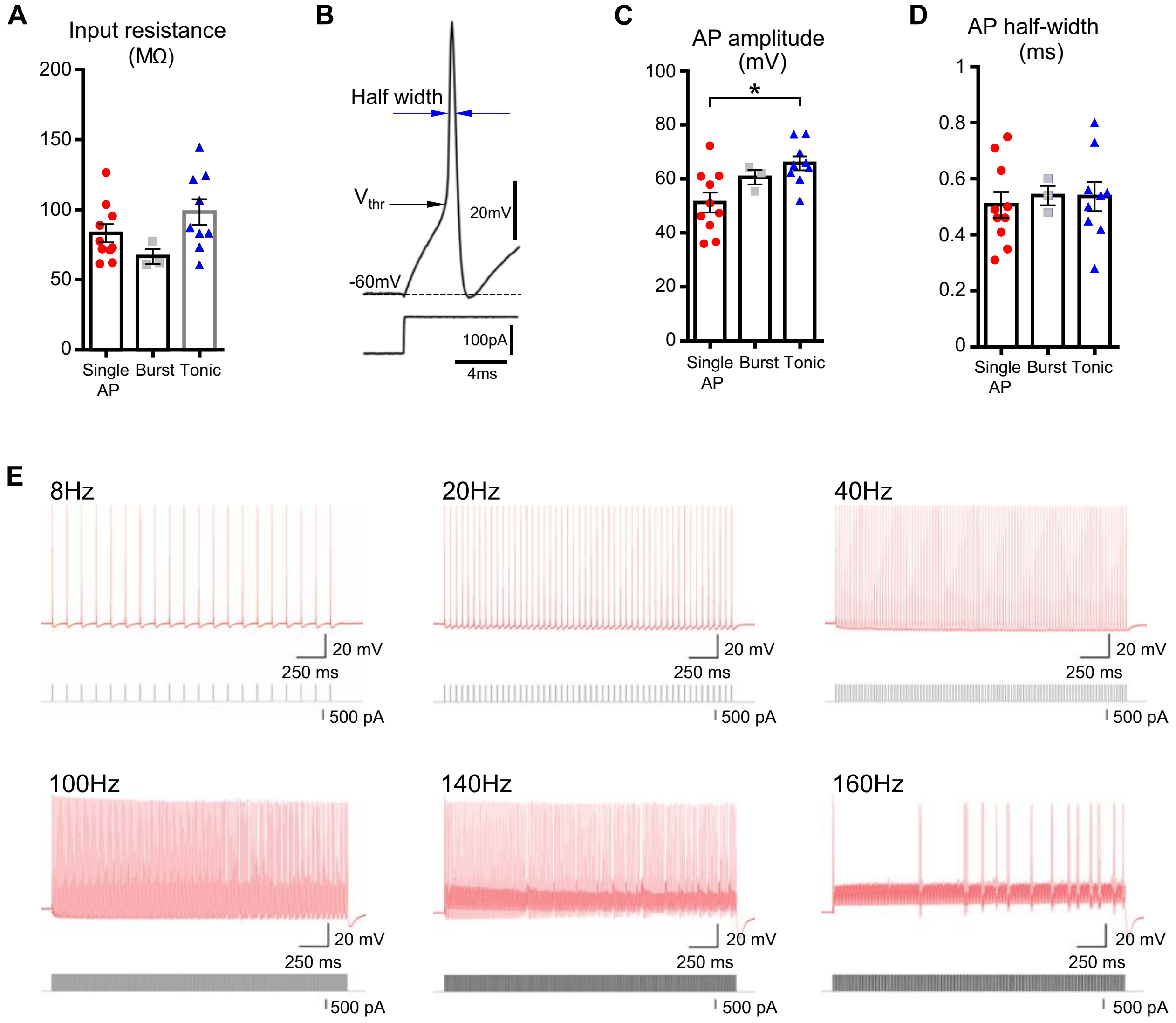

Supplemental Figure S12

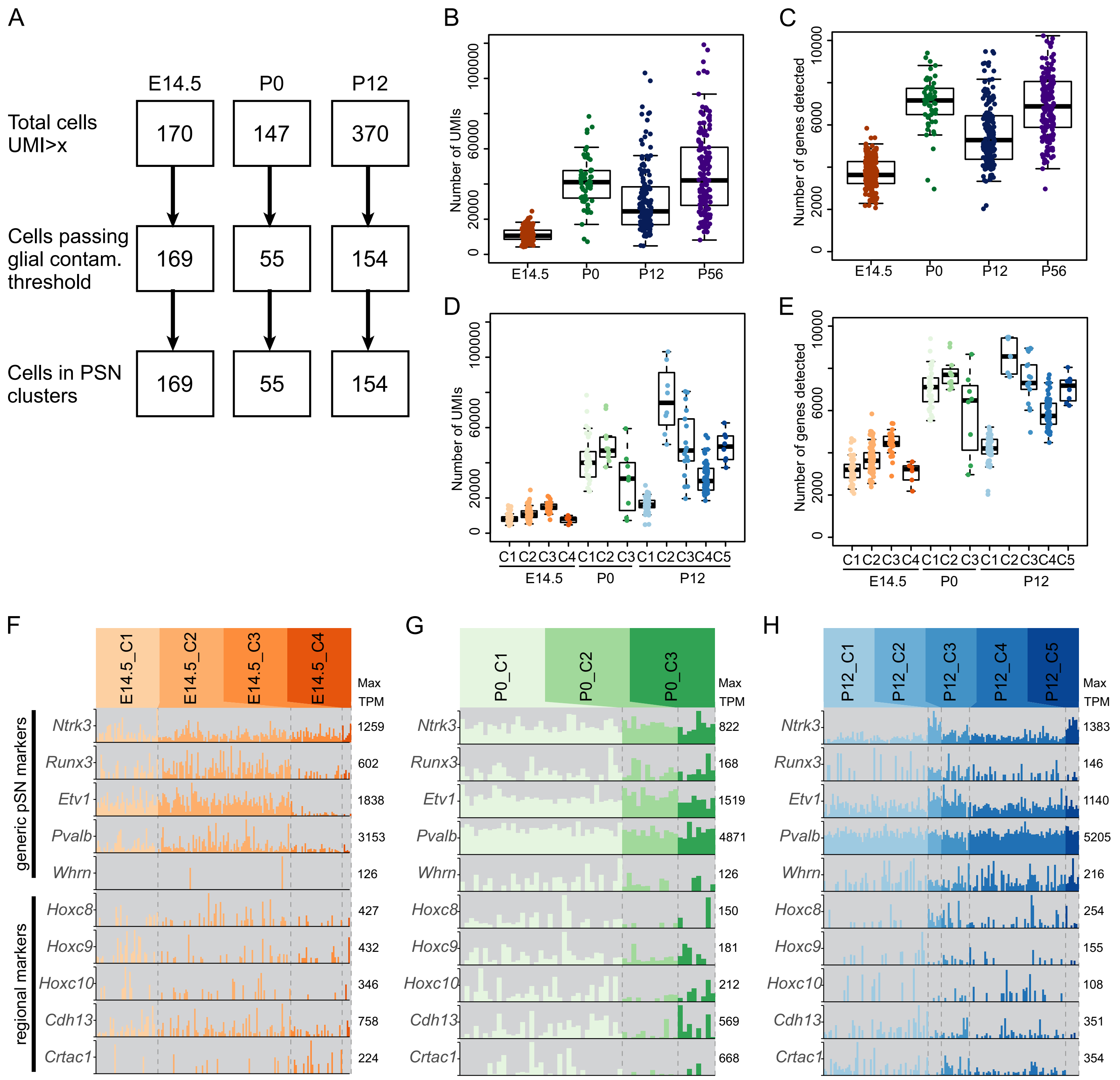

### Supplemental Figure 13

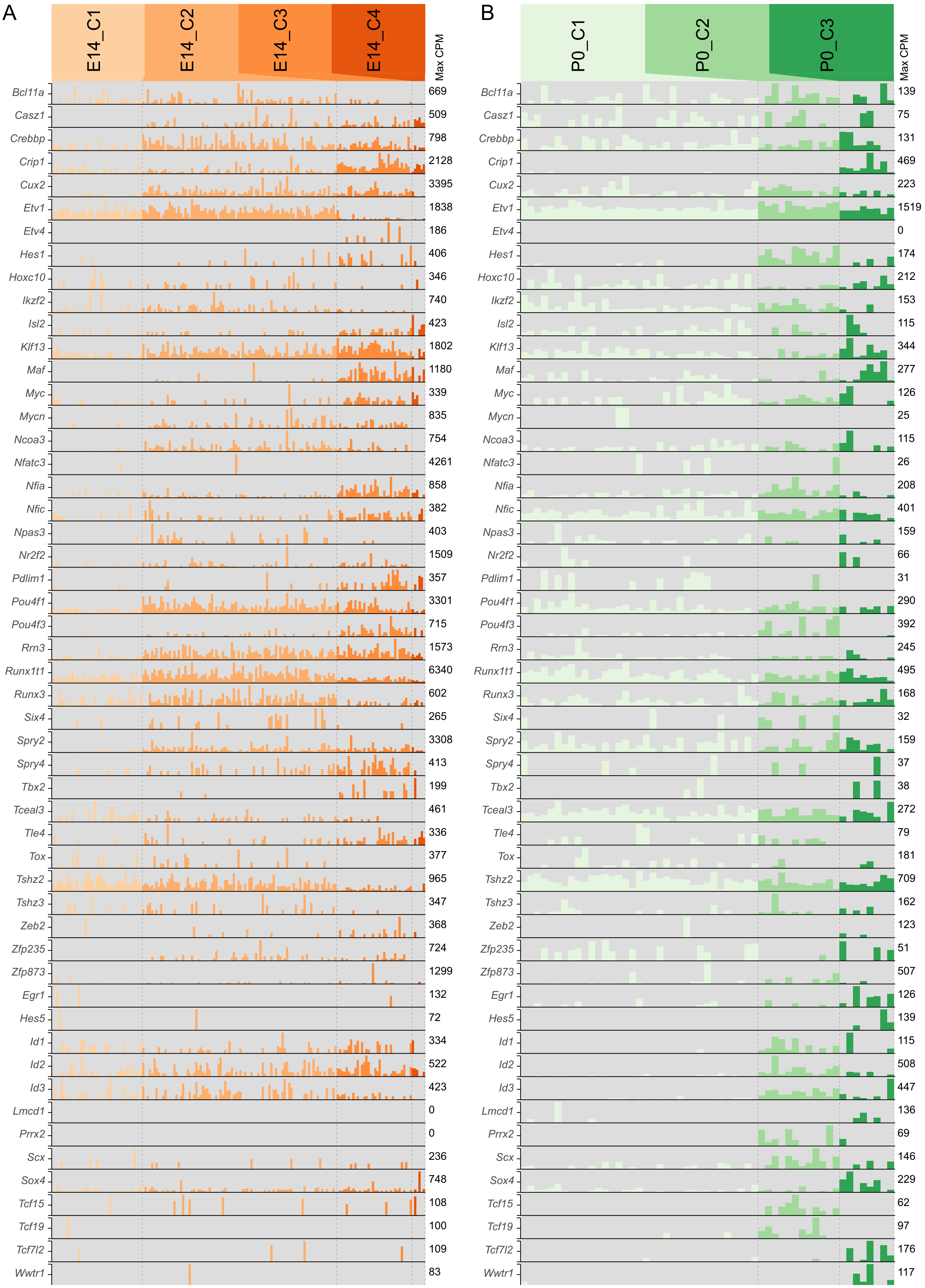

Supplemental Figure S14

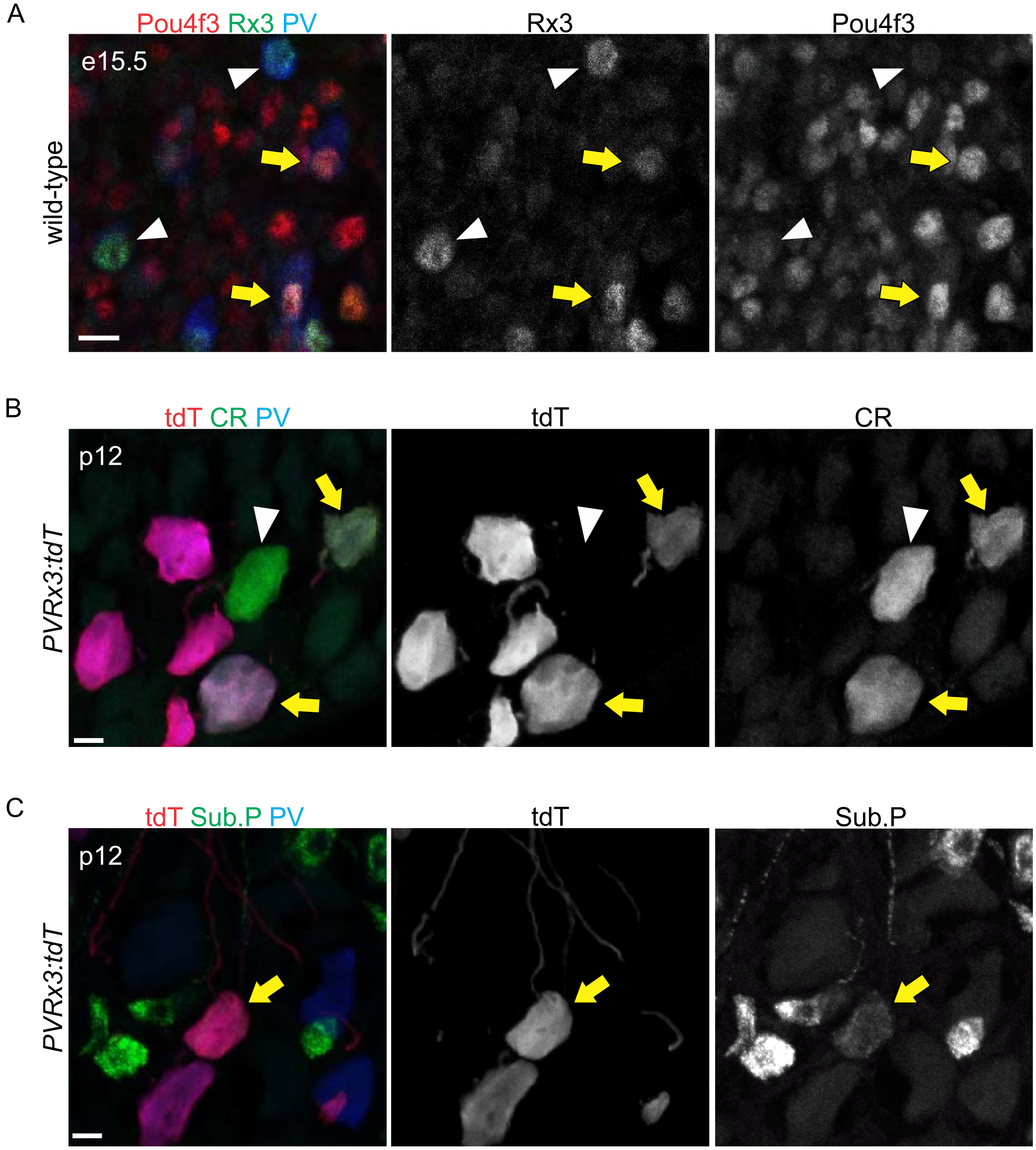
