## Supplemental Figure legends for "Molecular development of muscle spindle and Golgi tendon organ sensory afferents revealed by single proprioceptor transcriptome analysis"

**Supplemental Figures.**

**Figure S1. Phenotypic analysis of *Rx3:FlpO* mice.**

A. Diagram depicting the localization of the FlpO recombinase coding sequence within the *Runx3* genomic locus. Boxed areas indicate *Runx3* coding (in black) and non-coding (in gray) exons; the two Runx3 start sites are indicated (black arrows). An IRES-FlpO-neo cassette was inserted just after the Runx3 stop codon through homologous recombination in mouse embryonic stem cells. See methods for details.

B. Expression of *Runx3* (as assessed by RNA *is situ* hybridization) during early developmental stages indicates the gradual restriction of *Runx3* transcript to proprioceptive sensory neurons.

C. Illustration of genetic strategy to label Runx3^+^ DRG sensory neurons in *Rx3:FlpO;RCE* mice (*Rx3:GFP*).

D. Expression of Runx3, Islet1, and GFP in p2 lumbar DRG of *Rx3:GFP* mice. Boxed areas are enlarged in (*i, ii*). Corresponding images (*i’, ii’*) show expression of Islet1 (Isl1). Few Rx3^+^ neurons do not label with GFP (arrows), while GFP is observed in a substantial number of Rx3^off^Islet^+^ neurons (arrowheads).

E. Expression of Rx3, PV, and GFP in p2 L2 DRG of *Rx3:GFP* mice. Boxed areas are enlarged in (*i, ii*). Corresponding images (*i’, ii’*) show expression of GFP. Few Rx3^+^PV^+^ neurons do not label with GFP (arrow). While fragments of GFP^+^PV^+^ neurons (lacking a nucleus) can be detected (arrowheads), ‘full size’ (nucleated) GFP^+^PV^+^ neurons that lack Rx3 are not observed.

F. Percentage of Rx3^+^ neurons co-expressing GFP (of all Rx3^+^ neurons) in T10, L2 and L5 ganglia of p2 *Rx3:GFP* animals. Mean percentage for T10: 97.6 ± 0.5%, n = 2 DRG; L2: 91.2 ± 1.9%, n = 6 DRG; L5: 86.6 ± 1.8%, n = 2 DRG.

G. Percentage of Islet1^+^GFP^+^ neurons co-expressing Runx3 in T10, L2 and L5 ganglia of p2 *Rx3:GFP* animals. Mean percentage for T10: 63.1 ± 0.6%, n = 2 DRG; L2: 71.6 ± 3.4%, n = 2 DRG; L5: 72.6 ± 1.2%, n = 2 DRG. (When combined, 30.9 ± 2.1% of Islet1^+^GFP^+^ neurons lack expression of Rx3; n=6 DRG).

H. Percentage of Rx3^+^PV^+^ and Rx3^+^PV^off^ neurons (of all Rx3^+^ neurons) in L2 DRG of p2 *Rx3:GFP* animals. Mean percentage for Rx3^+^PV^+^: 83.4 ± 1.3%, n = 4 DRG; Rx3^+^PV^off^: 16.6 ± 1.3%, n = 4 DRG

I. Percentage of GFP^+^ neurons of all Rx3^+^PV^+^ or Rx3^+^PV^off^ neurons in L2 DRG of p2 *Rx3:GFP* animals. Mean percentage for GFP^+^Rx3^+^PV^+^: 95.7 ± 0.9%, n = 4 DRG; GFP^+^Rx3^+^PV^off^: 81.0 ± 5.8%, n = 4 DRG.

J. Expression of Runx3 and GFP in p12 spinal cord of *Rx3:GFP* animals. GFP is observed in central collaterals of pSNs as well as in a small number of cells that resemble microglia. No Rx3 or GFP expression was detected in spinal inter or motor neurons. Images *i*, and *ii* are taken from corresponding spinal areas indicated in schematic. Boxed area in *ii* is shown at higher magnification in *iii* and *iii’.*

K. Expression of Runx3 and GFP in p12 axial muscles of *Rx3:GFP* animals. GFP (and Rx3) is observed in small numbers of cells that resemble muscle satellite cells. Image taken from corresponding muscle area indicated in schematic of J. Boxed area is shown at higher magnification in *i* and *i’.*

Scale: 10 μm (B), or 20 μm (D, E, J, K).

**Figure S2. Phenotypic analysis of *PVRx3:tdT* mice.**

A. Expression of Runx3 and tdT in lumbar DRG of *PVRx3:tdT* animals. tdT^+^ neurons invariably express Rx3. Rx3^+^tdT^off^ neurons (arrowheads) correspond to PV^+^Rx3^+^ neurons in which Cre/Flp recombination failed, or to Rx3^+^PV^off^ neurons.

B. Percentage of tdT^+^Rx3^+^ neurons (of all Rx3^+^ neurons) in T10 and L5 ganglia of p28 *PVRx3:tdT* animals. Mean percentage for T10: 69.0 ± 4.4%, n = 17 sections; L5: 64.2 ± 2.9%, n = 83 sections.

C. Percentage of tdT^+^*PV*^+^ neurons (of all *PV*^+^ neurons) in rostral (L1-3) and caudal (L4-5) lumbar DRG of adult (≥p56) *PVRx3:tdT* animals. Mean percentage for L1-3: 86.9 ± 2.1%, n = 44 sections; L4-5: 73.9 ± 4.2%, n = 30 sections. Consistent with observations that RA mechanoreceptors are more prevalent at caudal lumber levels the percentage of tdT^+^*PV*^+^ neurons is higher at the L1-3 segmental level compared to L4-5 levels (p=0.004; Mann-Whitney U test).

D. Expression of *PV* and tdT in lumbar DRG of adult (≥p56) *PVRx3:tdT* animals. tdT^+^ neurons invariably express *PV*, but the level of *PV* transcript is highly variable (compare neurons indicated by arrowheads). *PV*^+^tdT^off^ neurons (asterisk) correspond to PV^+^Rx3^+^ neurons in which Cre/Flp recombination failed, or to PV^+^Rx3^off^ rapidly- adapting (RA) cutaneous mechanoreceptive neurons (e.g. Meissner, Pacinian, or Lanceolate afferents).

E. Expression of PV, Rx3, and tdT in lumbar DRG of p12 *PVRx3:tdT* mice. Some Rx3^+^tdT^+^ neurons express barely detectable levels of PV (dotted circles), consistent with observations in (D).

F. Expression of tdT in MS and GTO afferent collaterals in spinal cord of p12 *PVRx3:tdT* mice. Higher magnification of the dorsal horn in (*i, i’*) suggest that some tdT^+^ axons (arrow in *i’*) project to dorsal/intermediate spinal lamina.

Scale: 10 μm (A,C,E), 20 μm (G), or 50 μm (F).

**Figure S3: Analysis of cutaneous afferents in *PVRx3:tdT* mice.**

A. Illustration of genetic alleles enabling the selective labeling of PV^+^Rx3^+^ proprioceptive muscle afferents in DRG.

B. Expression of tdT, TuJ1 and S100 in forelimb glabrous skin of p2 *PVRx3:tdT* mice indicates association of tdT^+^ axons with TuJ1 labelled Merkel cells. Boxed area is shown at higher magnification in (*i-iv*).

C. Expression of tdT, Troma1, and S100 in forelimb digit nail of ≥p56 *PVRx3:tdT* mice. Boxed area is shown at higher magnification in (*i-ii*).

D. Expression of tdT, TuJ1 and S100 in forelimb glabrous skin of p2 *PVRx3:tdT* mice indicates absence of tdT^+^ axons from Meissner corpuscles (arrowheads). Boxed area is shown at higher magnification in (*i-ii*).

E. Expression of tdT and vGlut1 in the interosseous membrane of p2 *PVRx3:tdT* mice indicates absence of tdT^+^ axons from Pacinian corpuscles (arrowheads). Boxed area is shown at higher magnification in (*i-ii*).

F. Expression of tdT, Troma1 and S100 in (hairy) back skin of adult (≥p56) *PVRx3:tdT* mice. In contrast to glabrous fore paw skin, innervation of Troma1 Merkel cells in back (hairy) skin was never observed. Boxed area is shown at higher magnification in (*i-ii*).

G. Illustration of genomic alleles enabling the selective labeling of PV^+^Rx3^+^ muscle proprioceptor and PV^+^Rx3^off^ low-threshold cutaneous mechanoreceptors in DRG.

H. Expression of tdT, GFP and Troma1 in forelimb skin of ≥p28 *PVRx3:tdT*; *PV:GFP* mice showing tdT^+^GFP^+^ axons associated with Troma1^+^ Merkel cells (arrow). No tdT^+^ axons associate with GFP^+^ Meisner corpuscles (arrowhead). Boxed area is shown at higher magnification in (*i-iv*).

I. Expression of tdT, GFP and Troma1 in forelimb skin of ≥p28 *PVRx3:tdT*; *PV:GFP* mice simultaneously showing tdT^+^GFP^off^ axons associated with Troma1^+^ Merkel cells around a hair follicle (arrowhead), and tdT^off^GFP^+^ longitudinal Lanceolate endings (arrow). Boxed area is shown at higher magnification in (*i-iv*). We postulate that the mosaic activation of GFP in Merkel cell afferents may result from a) low *PV* transcript levels in these afferents, and from b) a higher Cre-mediated *loxP* recombination efficiency in the *Ai65:tdTomato* reporter when compared to the *PV:GFP* reporter.

Scale: 20 μm.

**Figure S4. Analysis of pSN single cell RNA sequencing data.**

A, B. Number of detected genes (all) (A) and number of detected transcription factors (TFs) (B) obtained for adult proprioceptors using either droplet-based (DB) or plate-based (PB) single cell sequencing platforms, using only the common genes in both data set transcriptome versions (to account for differences in total genes based on version differences). Detection was defined as a count value > 0. The number of detected genes (A) and transcription factors (B) per neuron is significantly higher for the plate-based sequencing method. (Ratio of mean genes detected in plate-seq vs. droplet (all genes): 1.81, DB mean (median) number of detected genes: 3304 (3240), PB mean (median) number of detected genes: 5989 (5968); p = 4.15e-55, Mann-Whitney test) (Ratio of mean TFs detected in plate-seq vs. droplet (TFs only): 1.90, DB mean (median) number of detected TFs: 231 (227), PB mean (median) number of detected TFs: 439 (436); p = 9.25e-52, Mann-Whitney test). DB-seq. data obtained from Sharma et al., 2020; PB-seq. data obtained from this study. Boxes show the median, 25^th^, and 75^th^ percentile, and whiskers extend to the most extreme data point less than 1.5 times the interquartile range.

C. Overview of the number of starting and filtered cells profiled using the plate-seq method (see Methods). Of the 450 initial plate wells, 208 cells passed the threshold for minimal cross-talk with non-neuronal transcripts, chiefly satellite cell genes such as Mbp (see Methods). Of these 208 cells, 173 were determined to be in the five clusters described in the main text, whereas the remainder fell into non-proprioceptor classes, or were in clusters with fewer than 10 cells.

D, E. Boxplots showing the number of (D) Unique Molecular Identifiers (UMIs) and (E) genes detected per cell, organized by cluster. Points for each individual cell are overlaid on the boxplots. Boxes indicate medians and 25^th^/75^th^ percentiles, and whiskers extend to the furthest point less than 1.5 standard deviations from the mean.

F. Barplots showing expression (counts per million, CPM) of previously reported pSN and regional markers, with individual cells arranged by cluster. The maximum CPM for each gene is reported on the right hand side.

G. Expression of previously identified pSN subset markers, *Heg1* (h), *Nxph1* (n), and *Pcdh8* (p), in lumbar DRG of adult (≥p56) *PVRx3:tdT* mice. Individual neurons (‘C1’-‘C5’; not part of larger images) serve as putative examples for the different classes of neurons uncovered through single cell RNAseq analysis.

H-J. Percentage of *Heg1^+^*tdT^+^ (F), *Nxph1^+^*tdT^+^ (G), and *Pcdh8^+^*tdT^+^ (H) neurons (of all tdT^+^ neurons) in rostral (L1-3) and caudal (L4-5) lumbar DRG of adult (≥p56) *PVRx3:tdT* animals. Mean percentage for *Heg1* (L1-3): 67.4 ± 4.4%, n = 14 sections; *Heg1* (L4-5): 59.8 ± 4.5%, n = 15 sections; *Pcdh8* (L1-3): 53.5 ± 2.3%, n = 50 sections; *Pcdh8* (L4-5): 54.4 ± 3.0%, n = 44 sections; *Nxph1* (L1-3): 62.3 ± 4.5%, n = 19 sections; *Nxph1* (L4-5): 53.5 ± 3.3%, n = 20 sections.

Scale: 10 μm.

**Figure S5. Validation of transcriptionally distinct pSN clusters.**

A, B. Expression of the cluster 5 transcripts *Pcdh17* and *Pcdh8* in ≥ p56 lumbar DRG of *PVRx3:tdT* mice. Images of individual neurons represent examples of observed transcript combinations other than those observed in the main image. (A) Pcdh17 (P_17_) transcript is generally only observed in tdT^+^ neurons with high levels of *Pcdh8* (p) transcript expression. (B) Percentage of *Pcdh17*^+^tdT^+^ neurons (of total tdT^+^) in ≥p56 rostral (L1-3) or caudal (L4-5) lumbar DRG of *PVRx3:tdT* animals. Median percentage ± S.E.M. at L1-3: 13.4 ± 3.0%, n=5 sections; L4-5: 23.8 ± 6.6%, n=7 sections.

C. Expression of cluster 1 transcripts *Hpse*, *Colq*, and *Agpat4* in *PV*^+^ neurons in ≥ p56 lumbar DRG of wild type mice. In (*i*) *Hpse* and *Colq* transcripts overlap and are largely restricted to *PV*^+^ neurons. In (*ii*) expression of *Agpat4* is observed in many neurons in DRG, but within *PV*^+^ neurons its expression typically overlaps with expression of *Colq*.

D. Expression of cluster 5 transcripts *Itga2*, *Pcdh17*, and *Chad* in *PV*^+^ neurons in ≥ p56 lumbar DRG of wild type mice. Expression of *Pcdh17* is observed in many neurons in DRG, but within *PV*^+^ neurons localizes to *Itga2*^+^ (*i*) or *Chad*^+^ (*ii*) PV neurons.

E. Expression of cluster 1 transcript *Colq*, in relation to the cluster 5 transcripts *Itga2*, *Chad,* and *Pcdh8*^+^ in ≥ p56 lumbar DRG of wild type mice.

Scale: 10 μm.

**Figure S6. Expression of Calretinin marks group Ia muscle spindle afferents.**

A. Colocalization of Calretinin (CR) with GFP in group Ia MS afferents in EDL muscle of p12 *PV:GFP* mice (top panels). In some MS afferents, expression of CR is not observed (bottom panels).

B. CR is not observed in GFP^+^ GTO afferents in EDL muscle in p12 *PV:GFP* mice.

C. Expression of CR and Calbindin 1 (CB) in motor neurons and motor axon endplates within neuromuscular junctions in p12 EDL muscle.

D, E. Genetic labeling of CR positive neurons in DRG using *Calb2:Cre* and *Mapt:lxp-STOP-lxp:GFP-iresNLZ*  alleles (hereafter *Calb2:GFP or Calb2:NLZ*). In (D) CR protein colocalizes with the nuclear localized βgalactosidase reporter. Consistent with the notion that CR labels a subset of proprioceptors, the *Calb2:NLZ* reporter is detected in only a small number of Rx3^+^ neurons (E).

F, G. Expression of GFP and vGlut1 in MS afferents in hind and fore limb muscle in p18 *Calb2:GFP* mice. In many muscles, GFP expression is localized to the vGlut1^+^ MS afferents that occupy the equatorial region of the spindle (F), but in some spindles GFP^+^ axons are not flanked by non-GFP^+^vGlut1^+^ afferents (a reflection of the heterogeneity with respect to muscle spindle anatomy/innervation patterns).

H, I. Consistent with expression of CR in motor neurons, in *Calb2:GFP* mice, expression of GFP can be observed in motor axons and motor end plates of alpha motor neurons (top panels), as well as in vGlut1^off^ gamma motor axons that innervate the MS (bottom panels).

Scale: 20 μm (C-G, I), and 50 μm (A, B, H)

**Figure S7. Expression of Pou4f3 in group Ib GTO afferents.**

(A-C) Genetic labeling of Pou4f3^+^ sensory neurons in *PV:Cre; Pou4f3:lxp-STOP-lxp:AP* (*PV/Pou4f3:AP*) animals.

(A) PV^+^ and Alkaline Phosphatase^+^ (PV^+^AP^+^) sensory neurons in p2 lumbar DRG. Higher magnifications in (*i, ii*) indicate different levels of AP labeling and may suggest a failure to maintain AP expression in some PV neurons.

(B) PV^+^AP^+^ sensory afferents in p2 gluteus, plantaris, and soleus muscle. In (*i*) AP^+^ GTO afferents and afferent terminals are readily observed, but AP^+^ MS afferents are no longer or just barely visible (arrows in *ii* and *iv*) at this stage.

(C) Spinal axon collaterals of PV^+^AP^+^ sensory afferents in p2 (*i, ii*) and ≥p56 (*iii, iv*) spinal cord at thoracic (*i, iii*) and lumbar (*ii, iv*) segmental levels. At p2, AP labeling is relatively abundant in all PV^+^ neurons and their spinal collaterals (*i, ii*). Consistent with the idea that group Ib GTO afferents primarily terminate on spinal interneurons located within intermediate spinal lamina, by p56, only traces of AP expression can be detected in ventrally projection neurons (*iii, iv*).

**Figure S8. Expression of Substance P in MS afferents.**

A. Expression of Substance P and GFP in p12 muscle in *Rx3:GFP* mice. An abundance of Tac1^+^ sensory terminals in muscle and MS sensory end organs appears to mask low Substance P expression levels in MS.

B. Expression of Substance P and tdT (*i*) and Rx3, PV, and tdT (*ii*) in p21 lumbar DRG of *Tac1:Cre; Ai14:tdTomato* (*Tac1:tdT*) mice. In (*i*) the discrepancy in tdT and Substance P expression provides evidence for a transient activation of *Tac1* in many DRG neurons at earlier developmental stages. Transiently expressing *Tac1* neurons include PV^+^Rx3^+^ proprioceptors (*ii*), providing a challenge to discern spindle afferents that temporally activated *Tac1* from those that maintain *Tac1* expression.

C. Expression of tdT and vGlut1 in >p28 muscle of *Tac1:Cre; Ai14:tdTomato* (*Tac1:tdT*) mice.

Scale: 20 (A, B), 50 (C) μm.

**Figure S9. Differential expression of biologically relevant transcripts across molecularly identified pSN subclasses.**

A. Go analyses of transcripts differentially expressed between molecular defined proprioceptor subtypes. Top-5 categories are shown only.

B, C. Barplots showing expression (counts per million, CPM) of cell surface molecules (B) and of transcriptional regulators (C). Genes belonging to each category were obtained from the Panther database, and filtered to include only those with significant differential expression between at least one pair of clusters (FDR p-value < 0.05) and expression in >= 50% of cells within at least one cluster.

**Figure S10. Distribution of ion-channels within muscle pSN subtypes.**

A, B. Barplots showing expression (counts per million, CPM) of calcium channel (A) and potassium channel (B) genes. Genes belonging to each category were obtained from the Panther database. Potassium channel genes were filtered to include only those genes with expression in at least one cell. Omitted K_v_ channels (with no appreciable expression in any pSN cluster) are *Kcna3, Kcna4, Kcna5, Kcna7, Kcnc2, Kcne1, Kcne1l, Kcnf1, Kcng3, Kcnh4, Kcnj1, Kcnj5, Kcnj8, Kcnj13, Kcnj15, Kcnk5, Kcnk6, Kcnk7, Kcnk9, Kcnk16, Kcnmb3, Kcnq1, Kcnq4, Kcns2, Kcnt2, and Kcnv2*.

**Figure S11. Action potential characteristics, passive membrane properties and ability of proprioceptive neurons to induce action potential firing at different frequencies.**

A. Measurements of input resistance for the three different types of proprioceptors observed in electrophysiological recordings.

B. The voltage threshold (V_thr_) and half width of an action potential (top) evoked on a proprioceptive neuron following current injection (bottom). Dotted line represents holding membrane potential at -60 mV.

C, D. Measurements of half-width of the action potential (C) and capacitance (D) for the three different types of proprioceptors observed in recordings.

(E) Representative voltage responses (top traces) following different frequencies of a small step of current injection (bottom) in the same neuron. Frequencies range from 8 Hz to 160 Hz.

**Figure S12. Analysis of developmental scRNAseq data sets.**

A. Overview of the number of starting and filtered cells profiled using the Plate-Seq method (see Methods) for three developmental time points. From the initial plate wells, cells were assessed for minimal cross-talk with non-neuronal transcripts, chiefly satellite cell genes such as Mbp (see Methods). Of the cells passing the contamination threshold, only those that ultimately belonged to clusters with >=5 cells were retained in the final step.

B, C. Boxplots showing the number of (B) Unique Molecular Identifiers (UMIs) and (C) genes detected per cell at each time point. Points for each individual cell are overlaid on the boxplots. Boxes indicate medians and 25^th^/75^th^ percentiles, and whiskers extend to the furthest point less than 1.5 standard deviations from the mean.

D, E. Boxplots showing the number of (D) Unique Molecular Identifiers (UMIs) and (E) genes detected per cell, subdivided by cluster. Points for each individual cell are overlaid on the boxplots. Boxes indicate medians and 25^th^/75^th^ percentiles, and whiskers extend to the furthest point less than 1.5 standard deviations from the mean.

F, H. Barplots showing expression (counts per million, CPM) of previously reported pSN and regional markers in the adult mouse, with individual cells arranged by cluster at each time point. The maximum CPM for each gene is reported on the right hand side.

**Figure S13. Transcription factor expression across e14.5 and p0 pSN molecular subtypes.**

A, B. Barplots showing expression (transcripts per million, CPM) of transcriptional regulators in cells from e14 (A) and P0 (B). Genes were obtained from the Panther database, and filtered to include only those with significant differential expression between at least one pair of clusters within a time point (FDR p-value < 0.05) and expression in at least 3 cells in one cluster.

**Figure S14. Expression pattern of newly identified MS and GTO afferent molecular markers in DRG.**

A. Expression of Pou4f3, Rx3, and PV in e15.5 lumbar DRG in wild type mice. Arrows indicate Pou4f3^+^PV^+^Rx3^+^ pSNs; Arrowheads indicate PV^+^Rx3^+^ pSNs with little or no Pou4f3 expression.

B. Expression of CR, PV and tdT in p12 lumbar DRG in *PVRx3:tdT* mice. Arrows indicate CR, PV^+^tdT^+^ pSNs; Arrowheads indicate Calretinin^+^tdT^off^ non-pSNs.

C. Expression of Sub. P, PV and tdT in p12 lumbar DRG in *PVRx3:tdT* mice. Arrow indicates SubP^+^PV^+^tdT^+^ pSN.

Scale 10 μm
